# Automatic Metadata Generation for Fish Specimen Image Collections

**DOI:** 10.1101/2021.10.04.463070

**Authors:** Joel Pepper, Jane Greenberg, Yasin Bakiş, Xiaojun Wang, Henry Bart, David Breen

## Abstract

Metadata are key descriptors of research data, particularly for researchers seeking to apply machine learning (ML) to the vast collections of digitized specimens. Unfortunately, the available metadata is often sparse and, at times, erroneous. Additionally, it is prohibitively expensive to address these limitations through traditional, manual means. This paper reports on research that applies machine-driven approaches to analyzing digitized fish images and extracting various important features from them. The digitized fish specimens are being analyzed as part of the Biology Guided Neural Networks (BGNN) initiative, which is developing a novel class of artificial neural networks using phylogenies and anatomy ontologies. Automatically generated metadata is crucial for identifying the high-quality images needed for the neural network’s predictive analytics. Methods that combine ML and image informatics techniques allow us to rapidly enrich the existing metadata associated with the 7,244 images from the Illinois Natural History Survey (INHS) used in our study. Results show we can accurately generate many key metadata properties relevant to the BGNN project, as well as general image quality metrics (e.g. brightness and contrast). Results also show that we can accurately generate bounding boxes and segmentation masks for fish, which are needed for subsequent machine learning analyses. The automatic process outperforms humans in terms of time and accuracy, and provides a novel solution for leveraging digitized specimens in ML. This research demonstrates the ability of computational methods to enhance the digital library services associated with the tens of thousands of digitized specimens stored in open-access repositories world-wide.

## I. Introduction

Over the last several decades advances in computing, imaging, and cyberinfrastructure have supported the growth of digital natural history collections, many of which contain specimen images [1]. Additionally, initiatives, such as the National Science Foundation’s Advancing Digitization of Biodiversity Collections (ADBC) program, have supported the digitization and curation of tens of thousands of biological specimens [2]. These digitized specimens are generally accessible through global, open-access repositories that support digital library services, such as browsing, search and retrieval, and preservation. The digitized renderings of these rich collections permit researchers, educators, students, and the general public to examine biological specimens on a previously unattainable scale. Moreover, the digitized instantiations present a pathway for making new scientific discoveries via the application of machine learning (ML).

Unfortunately, potential scientific advances are hindered by image quality problems and the lack of accurate and pertinent metadata associated with the image collections. Poor quality images (e.g. low contrast, inadequate lighting, out-of-focus or cluttered visual arrangements) are inadequate for automated image analysis by ML algorithms and lead to inferior computational results. In order to perform quantitative morphometric analysis of the specimens, the physical scale of the images 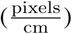 is needed; thus requiring the ability to identify and take measurements using rulers in the images. Many specimen collections do include Darwin Core metadata [3], detailing specimen taxon, geographic location, and several other specimen-related aspects. Additionally, some digitization efforts record technical metadata, detailing imaging specifications. While these types of metadata are helpful for a human examining several images at a time, they are insufficient for researchers seeking to apply computational methods to examine thousands of images to determine if, for example, a specific fish grows to different lengths in different habitats, or to study differences in the size of a particular anatomical feature, e.g. the size of a dorsal fin.

Since digital collections may each contain tens of thousands of images, manually producing image-related metadata for each digitized specimen is prohibitively expensive. Methods for automatically computing metadata are therefore needed to fully exploit biological image repositories for scientific discovery. As a step towards improving metadata in research specimen image collections, members of Drexel University’s Metadata Research Center are developing methods to automatically analyze fish images and extract a set of data features that provide important metadata about the digitized specimens. The research is being conducted as part of the Biology Guided Neural Networks (BGNN) project, which is developing a novel class of artificial neural networks that exploit machine readable and predictive knowledge associated with specimen images, phylogenies and anatomy ontologies. Using a combination of ML and image informatics techniques, we can accurately determine general image quality and metadata, such as fish quantity, location and orientation, and image scaling based on ruler identification and measurement. Image scaling allows us to compute quantitative features about the fish specimens, such as their length and area. In order to test and validate our methods, they have been applied to a set of 7, 244 images drawn from the Illinois Natural History Survey (INHS) Collection of fish specimens [4]. Figure 1 presents a typical image used in our study. The following section of the paper provides contextual background for this work, followed by the research goals and objectives, and a review of our research methods. Next, the results, along with discussion, are presented. The conclusion highlights key findings and identifies next steps.

**Fig. 1.**
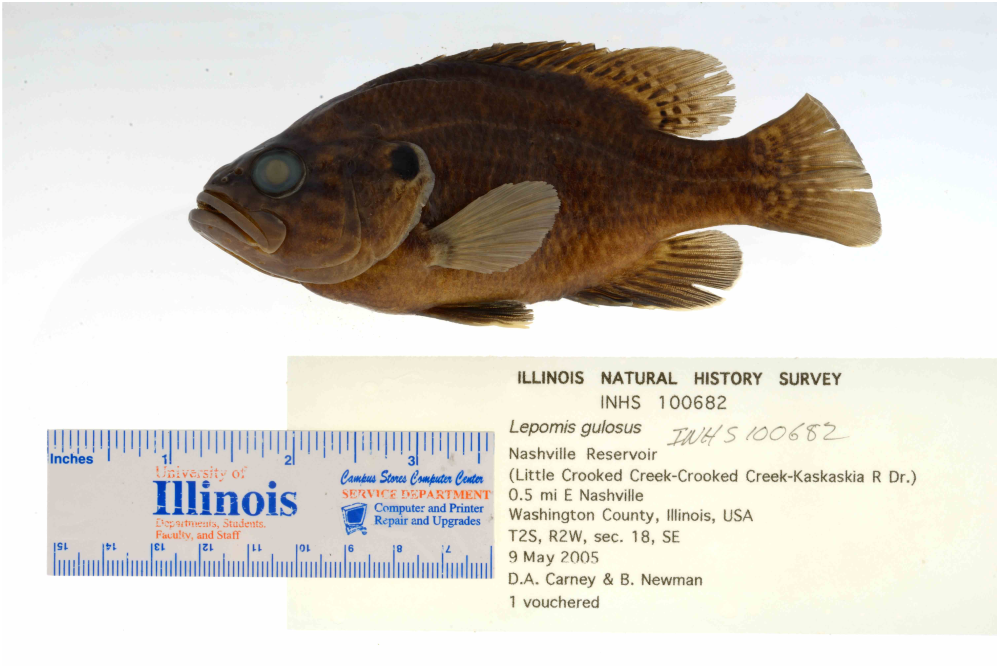
An image from the Illinois Natural History Survey (INHS) collection.

## II. Related Work

### A. Metadata for Natural History Image Collection

A number of different metadata standards have been applied to support the description and access of digital images of scientific specimens. The Darwin Core (DwC) [3], developed specifically to describe biological diversity data, is one of the most popular standards for such efforts. It is an extension of the Dublin Core’s DCMI Metadata Terms [5]. The Audubon Core [6], which supports the discoverability, dissemination, and use of data related to biological organisms (including 3– D digitized specimens), is a DwC extension that has become the popular metadata standard for biodiversity multimedia resources and collections. All of these descriptive standards and extensions include metadata properties for taxon, geographic location, and other important specimen content, and have been developed primarily from the perspective of a human curator. In other words, their application anticipates that a curator or data entry staff will manually generate metadata, drawn from acquisition logs or original specimen labels. The generated metadata associated with each digital rendering is generally sparse and prone to human error, placing limitations on a researcher seeking to apply ML to the image and the metadata for scientific research.

This limitation is magnified when trying to assess the actual quality of the digitized specimen. Descriptive-oriented standards support search and retrieval, and the biodiversity community has advocated for data fitness standards [7]. This point is also emphasized by Wieczorek et al. [8] in their report on the variety of DwC metadata extensions needed to meet growing community concerns and requirements, including data quality and fitness. Even so, metadata describing image quality is severely limited and generally missing. This point is addressed in detail by Leipzig et al. [9] and serves as the rationale for Tulane University’s effort to manually capture content for 22 metadata properties that characterize digitized specimen image quality. Their work is being conducted in connection with the larger BGNN initiative, and the difficulties encountered during the process underscore the need to explore automatic metadata generation methods.

### B. Automatic Metadata Generation

Advances in automatic metadata generation of both descriptive and technical metadata are relevant to the research presented in this paper. Automatic metadata generation of descriptive bibliographic data has been a research focus for close to 20 years [10]–[13]. Researchers have applied support vector machine (SVM) approaches [14], and associated networks to address sparse and incomplete metadata [15], and various successes are integrated into day-to-day workflows. Heidorn, et al. [16] demonstrated the use of optical character recognition (OCR) to extract specimen information from the original typed and often hand-annotated labels that are digitized along with herbarium collection holdings. The extracted information was encoded in the DwC metadata associated with the specimen’s digitized rendering. There has also been some success with extracting descriptive cartographic information from maps [17]. While descriptive metadata covers taxon, geographic location, and other important aspects, and may even record the image format; uses of automatic processes are still limited. More significantly, descriptive metadata does not sufficiently addressed data quality.

Technical metadata, such as camera settings and temporal information (date and time) are automatically generated during a digitization sequence, following standards such as Exchangeable image file format (Exif) [18]. The camera’s technical metadata is automatically captured and inserted into digital image files at the time of acquisition. Some of this metadata may be useful when selecting a ML sample. A researcher may desire images with specific properties, such as being captured chiefly with a certain aperture setting. Even so, the majority of automatically registered technical metadata associated with digitized specimens is also insufficient for computational research, leaving researchers to rely on manually generated descriptive metadata, which itself is sparse and prone to human error. Fish image analysis research, as reviewed below, demonstrates the potential of automated computational methods to address current metadata shortcomings and needs specific to the selection of high-quality digitized specimen images for the application of ML.

### C. Fish Image Analysis

Image analysis has been utilized to examine and process images of fish for well over two decades [19], [20]. It is an important application of technology for marine science, in the study of aquatic species, habitats and ecosystems, and for the seafood industry, in the development of automated fish sorting and grading systems, as well as fisheries management. Many of these computational analyses focus on the recognition and classification of the fish present in an image. The computational methods employed for fish image analysis have followed the general trends in the AI field. Hu et al. [21] presented a method of classifying species of fish based on color and texture features and a multi-class support vector machine (MSVM) [22]. Li and Hong [23] computed eleven shape and color features from fish images and derived a linear model that could discriminate between four different fishes. Rodrigues et al. [24] explored several combinations of feature extractions, input classifiers and clustering algorithms to produce a method that could distinguish between 10 different types of fish with 92% accuracy. Salman et al. [25] employed a deep Convolution Neural Networks (CNN) [26] together with classification based on K-Nearest Neighbor and Support Vector Machines trained on the features extracted by the CNN. They achieved 90% accuracy when identifying 15 different fish species in challenging underwater digital images. Utilizing texture, anchor points, and statistical measurements, Alsmadi et al. [27] implemented fish classification through a metaheuristic algorithm known as the Memetic Algorithm. They were able to classify 24 fish families with 90% accuracy. Iqbal et al. [28] used a modified AlexNet [29] model to classify six different fish species with 90% accuracy.

Especially in industrial settings, it is necessary to automatically detect the orientation, length and weight of fish during handling and processing. In some instances fish in the images need to be computationally straightened before further processing can be attempted [30]. Balaban et al. [31] demonstrated that image analysis and data fitting may be used to predict the weight of salmons with high accuracy. Hao et al. [32] provide an excellent review of fish measurement efforts that utilize machine vision. Azarmdel et al. [33] developed a system capable of determining the orientation of a trout and segmenting its fins, which are used as cutting points, with an accuracy over 99%.

The research reviewed above demonstrates the application of image feature extraction and machine learning algorithms to fish images; although researchers have not applied these approaches to the numerous collections of digitized specimens accessible in open repositories. Our research addresses this need by applying ML and informatics techniques to extract key metadata properties from the images. The availability of general and powerful off-the-shelf ML tools make the usage of previous special-purpose techniques unnecessary.

## III. Goals and Objectives

Digitized specimens accessible in open-access repositories provide a rich, extensive data source for ML and scientific discovery. These resources, however, remain largely untapped due to image quality issues and metadata limitations. The overall goal of our work addresses this need by developing a computational alternative to the current manual metadata generation process, which is prohibitively costly both in terms of labor and time. Additionally, our methods collectively provide a novel and general approach to computing higher-level metadata that will support scientific inquiry based on the analysis of specimen image collections.

Our four key objectives are to:

1) Explore use of Facebook AI Research’s `detectron` tool. Specific aims are to use `detectron` to identify study-specific objects.
2) Investigate image processing at the pixel level. Pilot testing found that `detectron` undersegmented the detected objects with tightly enclosing bounding boxes. We will determine if pixel analysis methods commonly found in image informatics may produce more accurate bounding boxes and object masks. The specific aims of this objective are to:
  a) Identify the appropriate threshold value for a more accurate mask.
  b) Remove noise to produce a single, solid mask.
  c) Compute a more accurate bounding box from the updated mask.
  d) Automatically determine when modified methods fail and `detectron` values should be used as is.
3) Compute a number of high-level metadata properties from the detected objects and image quality metrics.
4) Compare computed metadata properties with manually generated properties when possible to assess the accuracy and effectiveness of automated methods.

The automated metadata generation methods for our project were developed to work on a specific set of images from the INHS Fish Collection [4]. Most of these images have been configured, produced and acquired with a standard procedure. The images used for our study contain one fish placed on a bright, white background and contain an information tag and the same ruler. See Figure 1 for an example image from the collection. While training and focusing our system on images with very similar compositions and visual properties may limit its immediate applicability, our efforts demonstrate the potential that ML and image informatics techniques have for automatically generating metadata for biological specimen image collections in general.

## IV. Methods

Our process for metadata generation can be divided into three steps: 1) object detection with Facebook’s Detectron2 ML library (referred to as `detectron`), 2) image processing at the pixel level, and 3) calculations on the results of the previous steps to determine higher level metadata properties.

### A. Detectron

A prerequisite task to performing any advanced metadata property generation is finding the specimens (and other relevant objects) within the collection images. Object detection has been a broadly active field of study in recent years [35], and has resulted in a number of well-tested, purpose-built architectures. We elected to use Facebook AI Research’s (FAIR) `detectron` tool [34], and specifically its implementation of the Mask R-CNN architecture [36], for object detection in our project, given its many flexible and robust capabilities. Most importantly, following a review of the literature and available tools, we determined that there were no other machine learning packages that returned pixel by pixel masks over detected objects in a comparable fashion.

`detectron` is built on `pytorch` [37] and provides a relatively straightforward method for training on COCO [38] format datasets. It is able to handle any number of object classes, and can classify an arbitrary number of objects within a given image. We chose `detectron` for its relative ease of use compared to lower level libraries, and its implementation of powerful architectures developed by FAIR. For our project, we use it to identify five object classes: fish, fish eyes, rulers, and the numbers 2 and 3 on rulers, as shown in Figure 2. Objects with a 30% confidence score or higher are maintained for analysis.

**Fig. 2.**
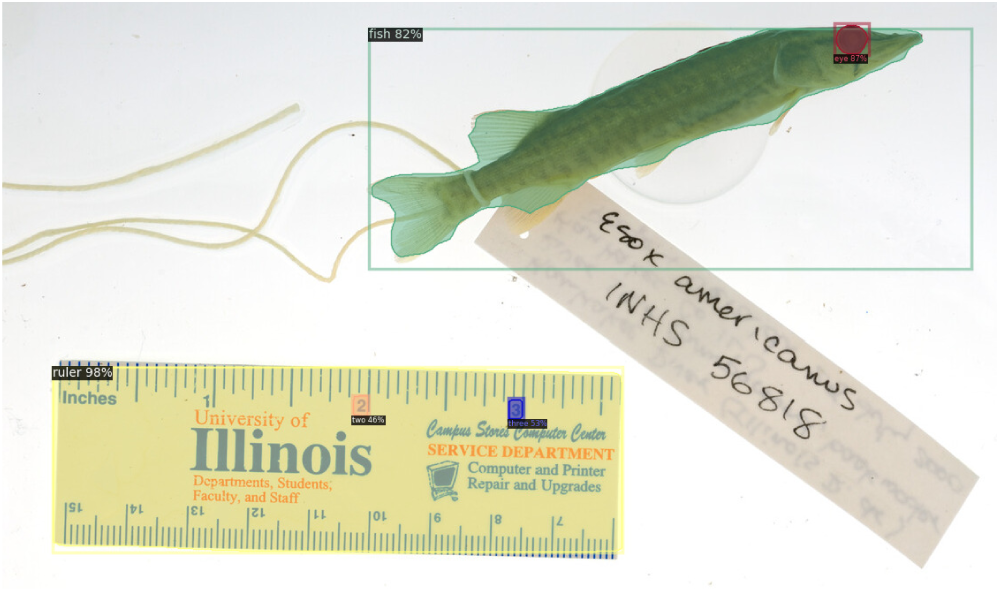
Initial object detection on a specimen image using Detectron2 [34].

Table I lists the number of instances for each class used in our training dataset. All of the training data was labeled by hand using `makesense.ai` [39] on images from the INHS Fish Collection [4]. Using `detectron`’s default training scheme, the model was trained for 100, 000 epochs. All instance types were included in a single object detection model.

**TABLE I.** Training dataset

| Class | Number of Instances |
| --- | --- |
| Fish | 297 |
| Ruler | 1496 |
| Eye | 456 |
| Two | 100 |
| Three | 100 |

### B. Pixel Analysis

The masks and bounding boxes produced by `detectron` are generally quite good, although they almost never completely or tightly enclose the detected objects. This is problematic for the detected fish objects in our analyzed images, where the most accurate segmentation is desired. The mask may include additional background as part of the fish, or the bounding box may clip away part(s) of the fish. To solve these shortcomings, we utilize pixel analysis methods commonly found in image informatics to produce more accurate object masks and bounding boxes.

#### 1) Threshold Adjustment

The first calculation in the pixel analysis process determines the cutoff intensity between what constitutes the foreground (i.e. the fish) and background of the image. Initially, the calculation is based on the bounding box and mask generated by `detectron`. Specimen images are read in as gray scale, and pixels in the image are treated as unsigned integers between 0 and 255. Otsu thresholding [40], a technique that maximizes the variance between the foreground and background intensities, is used to compute an initial cutoff value between foreground and background. While the Otsu value occasionally generates an accurate mask as is, usually the contrast between foreground and background is low and much of the lighter parts of the fish (such as its tail fin) are marked as background.

To overcome this improper segmentation, the threshold value should be either adjusted up or down, depending on whether the background is lighter or darker than the fish. For our current dataset, the background is always lighter (i.e. closer to 255), so the threshold value needs to be scaled up to include more of the foreground image. For optimal results the scaling should be dependent on the contrast between the background and foreground, which can be affected by the level of pigmentation of the fish. We found that an improved threshold value can be computed as the halfway point between the Otsu threshold value and the mean of the background intensities. This adjusted threshold value usually produced an acceptable balance between capturing most of the fish’s fins, without also masking parts of the background.

#### 2) Consolidating the Foreground

While thresholding has the potential to generate better masks than a neural network (when provided an initial approximate bounding box), it also introduces considerable noise. Single or small groups of errant pixels can be marked as foreground depending on the consistency of the background, and interior pixels of the fish (especially around the fins) can be marked as background. To be useful for generating an accurate bounding box and for subsequent computational analysis, the mask must consist of one single “blob” over the fish, i.e. containing no holes, and no other pixels disconnected from this blob can be marked as foreground.

To accomplish this, we apply an iterative process of flood filling from all the foreground pixels in the image until a blob is generated that is large enough to constitute the fish. This leads to another metaparameter, but using greater than 10% of the current bounding box has masked the specimen in all observed cases. Once the fish’s blob is found, noise then needs to be removed. This is done by flood filling from each of the corners of the bounding box, where the specimen is not present (all four corners in the overwhelming majority of cases), then taking the inverse of the result. The fish mask is excluded from these corner flood fills, so this process removes all noise from both the background and foreground of the image, leaving only a single mask over the fish itself.

#### 3) Adjusting the Bounding Box

With an accurate mask generated, it is then necessary to check whether the bounding box needs to be expanded or shrunk along any of its edges. Expansion is done first, by checking whether any edge intersects with any of the foreground mask pixels. If one does, it is expanded out by 1 pixel. If any edges are expanded, the whole process of masking and expansion is repeated to account for any changes in average intensities. Once no edges contain foreground pixels, the bounding box is then shrunk. Each edge is contacted by one pixel until it contains one or more foreground pixels. Once the shrinkage step is accomplished, the final mask and bounding box have been generated.

#### 4) Fallback

The pixel analysis process occasionally fails, e.g. when flood-filling does not produce a large enough blob or the bounding box adjustment does not terminate. This can occur if certain flood fill operations behave unexpectedly, or if the image is too washed out or otherwise atypical for the thresholding process to work correctly. In the event this happens, the original mask and bounding box generated by `detectron` is used for metadata generation.

### C. Metadata Generation

The following metadata properties are generated from the methods described above: `has_fish`, `fish_count`, `has_ruler`, `ruler_bbox`, `background.{mean,std}`, `foreground.{mean,std}`, `bbox`, `mask`, `score`, and `has_eye`. The remaining properties are computed as described below, with all the metadata properties listed in Table II.

**TABLE II.** Metadata properties (* indicates higher order derived properties)

| Property | Association | Type | Explanation |
| --- | --- | --- | --- |
| has_fish | Overall Image | Boolean | Whether a fish was found in the image. |
| fish_count | Overall Image | Integer | The quantity of fish present. |
| has_ruler | Overall Image | Boolean | Whether a ruler was found in the image. |
| ruler_bbox | Overall Image | 4 Tuple | The bounding box of the ruler (if found). |
| scale* | Overall Image | Float | The scale of the image in $\frac{\text{pixels}}{\text{cm}}$. |
| bbox | Per Fish | 4 Tuple | The top left and bottom right coordinates of the bounding box for a fish. |
| background.mean | Per Fish | Float | The mean intensity of the background within a given fish's bounding box. |
| background.std | Per Fish | Float | The standard deviation of the background within a given fish's bounding box. |
| foreground.mean | Per Fish | Float | The mean intensity of the foreground within a given fish's bounding box. |
| foreground.std | Per Fish | Float | The standard deviation of the foreground within a given fish's bounding box. |
| contrast* | Per Fish | Float | The contrast between foreground and background intensities within a given fish's bounding box. |
| centroid | Per Fish | 4 Tuple | The centroid of a given fish's bitmask. |
| primary_axis* | Per Fish | 2D Vector | The unit length primary axis (eigenvector) for the bitmask of a given fish. |
| clock_value* | Per Fish | Integer | Fish's primary axis converted into an integer "clock value" between 1 and 12. |
| length* | Per Fish | Float | The length of a fish in centimeters. |
| mask | Per Fish | 2D Matrix | The bitmask of a fish in 0's and 1's. |
| pixel_analysis_failed | Per Fish | Boolean | Whether the pixel analysis process failed for a given fish. If true, detectron's mask and bounding box were used for metadata generation. |
| score | Per Fish | Float | The percent confidence score output by detectron for a given fish. |
| has_eye | Per Fish | Boolean | Whether an eye was found for a given fish. |
| eye_center | Per Fish | 2 Tuple | The centroid of a fish's eye. |
| side* | Per Fish | String | The side (i.e. 'left' or 'right') of the fish that is facing the camera (dependent on finding its eye). |

#### 1) Contrast

The contrast between the intensities of the foreground and background pixels is computed as `background.mean` - `foreground.mean`.

#### 2) centroid and eye center

Centroids are provided for the masks and bounding boxes generated by `detectron`, and since we do not recalculate the mask of fish eyes we can use that value directly for `eye_center`.

Since we recalculate the mask of the fish, its centroid must be recalculated as well. This can be done via

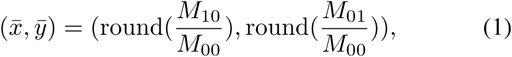

where *M*_00_ is the pixel area of the fish’s blob, *M*_10_ is the sum of all the *x* values of blob pixels, and *M*_01_ is the sum of all the *y* values of blob pixels.

#### 3) Side

Determining which `side` of the fish is visible is predicated on finding its eye. If an eye is found, the sign of the *x* component of the vector from the centroid of the fish to the centroid of the eye specifies which side is up: negative for left and positive for right. This assumes the fish was photographed vertically (i.e. dorsal fin on top), which is essentially always the case for all image collections our group has worked on.

#### 4) primary _axis and clock _value

The `primary_axis` of a fish can be calculated by taking the covariance of its blob in *x* and *y*, which yields its principle eigenvector. The eigenvector can be directly assigned to the property. If an eye is present, we ensure that `primary_axis` points in the direction of the eye relative to the fish’s centroid.

Our team encoded this information as a “clock value” between 1 and 12 when manually recording it. To convert `principal_axis` to `clock_value`, the sign of *x* and *y* on the principal axis are used to determine which Cartesian quadrant the fish angles into relative to its centroid. Depending on the quadrant, we dot product the principal axis with either [−1, 0], [0,−1], [1, 0] or [0, 1], which correspond to 9, 6, 3 and “0” o’clock respectively. The resulting radian value is then converted to a polar displacement in clock value space, and added to the comparative clock value used in the dot product. This gives the fish’s clock value from 0 to 11.9. Before recording `clock_value` in the output, the value is rounded to the nearest integer, with a 0 final result replaced with 12.

#### 5) scale and length

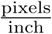 can be calculated by measuring the distance in pixels between the digits 2 and 3 (a 1 inch separation) found on the ruler by `detectron`. Converting this to 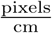 gives the `scale` metadata property as reported in the output.

For the fish `length` property, it is necessary to determine the furthest points from the centroid of the fish in each direction along its major axis. Since fish are normally measured in a straight line from their snout down the middle of their trunk, every pixel of the fish blob is projected onto the fish’s major axis (as a line through its centroid). The projection is done by finding the closest point on the centroid–principal axis line from the pixel’s location. After processing every pixel in the fish blob, computing the distance between the two furthest projected points gives the length of the fish in pixels. Multiplying this distance by `scale` gives the fish `length` in centimeters.

## V. Results

Technicians employed by Tulane University have manually generated the 22 metadata properties deemed crucial to the overall BGNN project [9] for a large number of INHS images. 20, 699 total entries were created by 13 technicians that spanned 8, 398 unique images, of which 7, 244 were both not part of the `detectron` training set and met our current admissibility criteria for `detectron` and pixel processing. We ran the metadata extraction program on these 7, 244 images. For the properties of image scale, fish length, and fish bounding boxes (properties not manually generated), a random sample of 100 specimens from the set of 7, 244 were analyzed by hand for comparison.

Our automated process currently generates 6 of the 22 core metadata properties: `if_fish` (`has_fish`), `fish_number` (`fish_count`), `if_ruler` (`has_ruler`), `specimen_angled` (`clock_value`), `specimen_view` (`side`), and `brightness` (`foreground.mean`). In addition, our approach also calculates contrast, bounding boxes and fish lengths in centimeters.

### A. Fish Detection

All images in the INHS dataset contain exactly one fish. For 7, 209 of the specimen images, one fish was detected, a 99.5% correct rate. For 25 of the images, 2 fish were detected, 3 fish were detected for 3 images, and for 7 images no fish were found. The 7 fish that were not detected were quite small. This type of specimen is currently lacking from the training set. See the top image in Figure 3 for an example. In the case of greater than 1 fish, 9 of the 28 contained tags that overlapped the fish and were themselves labeled as a second fish. Of the remaining 17, `detectron` erroneously labeled the fish as two separate fish objects, or labeled a subsection of the fish a second time. Fish that were double labeled were generally quite large and/or dissimilar from the fish found in the training set, such as the bottom image in Figure 3.

**Fig. 3.**
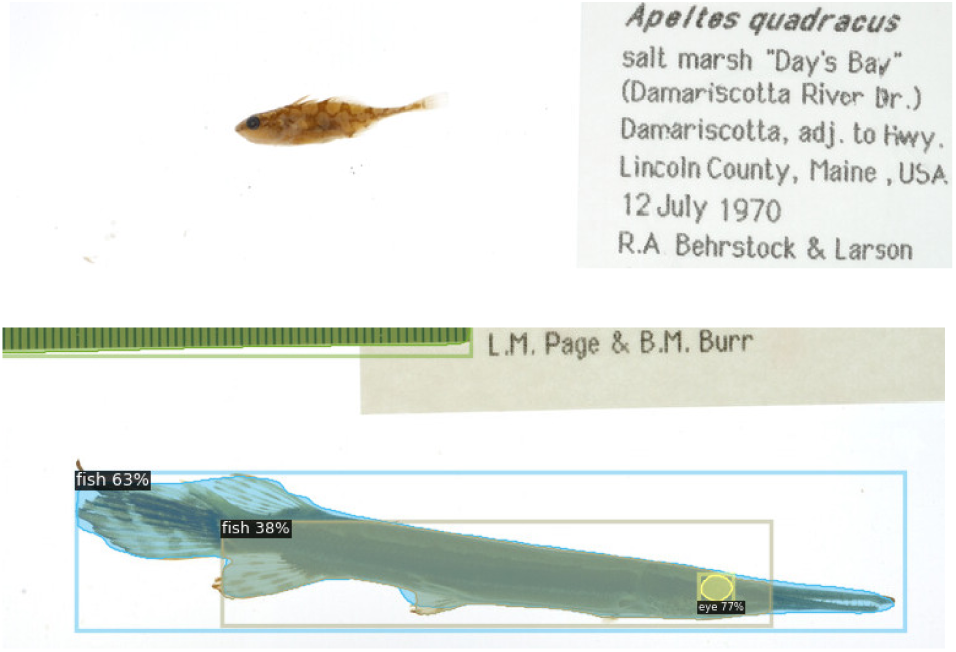
A fish that was not detected (top) and a fish that was detected twice (bottom).

### B. Ruler Detection

For all but 2 of the 7, 244 images `detectron` was able to find the ruler, a nearly perfect correct rate. In 56 of the images, the ruler itself was found, but the numbers “2” and/or “3” on the ruler were not. Therefore, a scale calculation could not be performed, producing a 99.2% success rate for the `scale` computation.

Images where one of these objects were not detected generally had some form of coloration issue. They were either washed out, very dark or yellow in hue. See Figure 4 for two such examples. Some of the rulers for which “2” and/or “3” were not detected were particularly scratched and damaged. Only “3” was missed in 45 of the 56 cases, only “2” was missed in 2 cases, and both were missed in 9 cases. This may indicate that more training samples for “3” are required. Many of the rulers where both numerals were missed were particularly small within the image, which again may be solvable through expanding the training dataset.

**Fig. 4.**
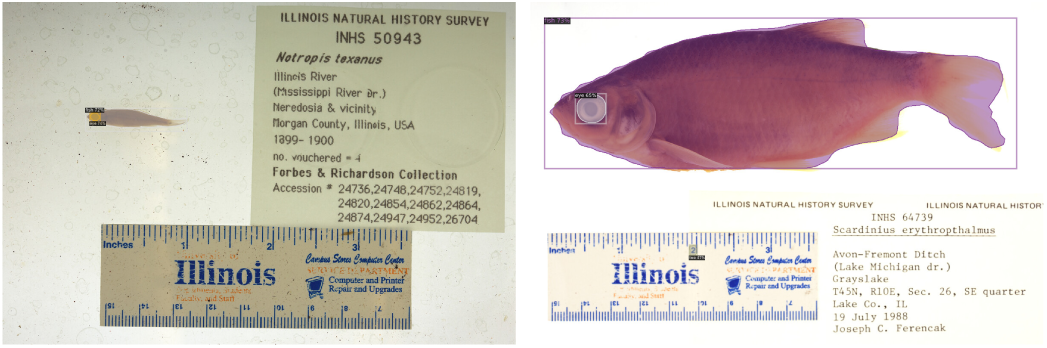
The two images where the ruler was not found. The left image exhibits a yellow hue, and the right is quite washed out with poor contrast.

### C. Side Detection

`detectron` was unable to find a fish eye in 246 of the images. These eyes were generally extremely dark, small, or looked nothing like those found in the training set. Of the remaining 6, 998 images, the correct side (`left` or `right`) was detected in all but 6 cases, producing a 96.5% correct rate.

For these 6 incorrect cases, a spot on the wrong side of the fish was labeled as the most likely eye within the bounding box of the fish. Figure 5 presents one such example. There were an additional 17 images for which the automated process generated a result that did not match the manually created data. For these remaining cases the manual data was incorrect, giving the automated system an error rate 2.8 times lower than the human error rate. This result highlights the additional utility of automated methods to double-check and verify manually generated metadata.

**Fig. 5.**
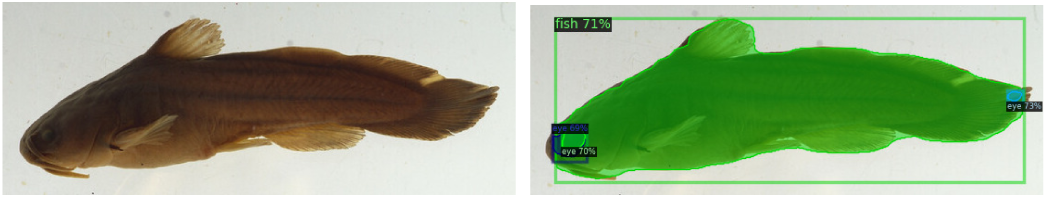
A fish for which a splotch on its tail fin was labeled the most likely eye.

### D. Clock Value

Clock position values were successfully generated for 6, 991 of the images. Of those, all but 8 were within ±1 of the correct result, our definition of a correct/acceptable result, making the correct rate for this computation 96.4%.

Of the specimens for which clock values were generated, 33 did not match the manually created data (within a tolerance of ±1). For 25 of those, the manually generated data was incorrect, giving the automated process a 3.1 times lower error rate. Of the 8 that were computationally classified incorrectly, two specimens were quite curved making it difficult to assign a clear angle value, as seen in Figure 6. The other 6 were values of “3” instead of “9” or vice versa, which resulted from a mislabeled eye as discussed in the previous section.

**Fig. 6.**
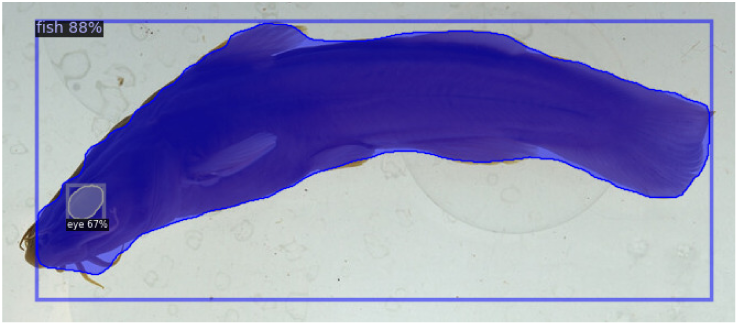
An example of a heavily curved specimen.

### E. Image Contrast

The contrast of an image is an important image property needed for the analyses of the BGNN study. Ideally, for INHS images the fish should be well lit and clearly displayed, and the background should also be as light and white as possible. The difference between the mean intensity of the foreground (i.e. the pixels of the fish) and the mean intensity of the background within the fish’s bounding box was computed for all 7, 244 images. The overall mean of the differences is 144.3, with a standard deviation of 15.8. Images on the low end of the distribution exhibit poor foreground–background contrast, and images on the high end likely contain poorly lit specimens. Images are considered to have “low” and “high” contrast if their background-foreground difference is greater than one standard deviation away from the mean; otherwise they are classified as “medium”. Examples of each type of image (low, medium, and high contrast) can be seen in Figure 7. The left image has a contrast of 103 (low), the middle image 144 (medium), and the right 186 (high).

**Fig. 7.**
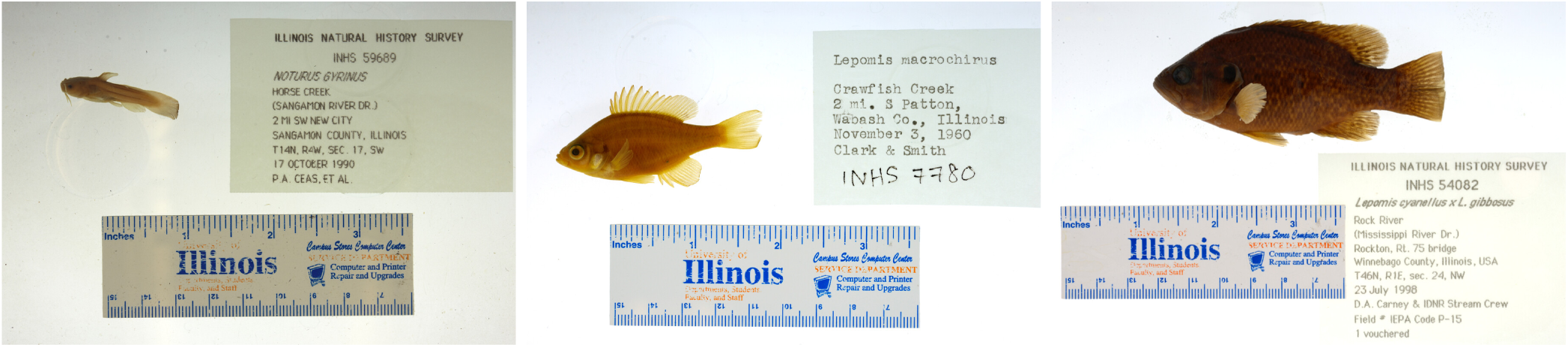
Examples of low, medium and high contrast fish images.

### F. Brightness

Specimen brightness is one of the 22 hand-recorded meta-data properties. It is encoded as `dark`, `normal` or `bright`. These values correspond to the mean foreground intensity computed by the automated system. The mean and standard deviation of the foreground intensities were computed for the images in the three manually specified classes. Table III contains the resulting values and shows that automated intensities values provide objective measures that may be used to break images into groups that correlate with manually generated brightness classifications. The mean of `foreground.mean` over all 7,244 images is 89.5, with a standard deviation of 16.6. Given that human perception of brightness is quite subjective and highly variable, we found it problematic to use specific threshold values to define `dark`, `normal` or `bright` specimens in concordance with the manually generated metadata. The technicians showed little consistency when classifying `normal` versus `bright` specimens. We did find that an 81.2% accuracy rate could be achieved by classifying `dark` images as those with a `foreground.mean` of 75 or less; thus demonstrating that this computed metadata property does offer some value when assessing image quality.

**TABLE III.** foreground.mean statistics for the three Brightness classes

| Class | Mean Intensity | Standard Deviation | Instances |
| --- | --- | --- | --- |
| Dark | 75.2 | 13.5 | 1800 |
| Normal | 93.6 | 14.8 | 5186 |
| Bright | 108.4 | 15.2 | 242 |

### G. Mask and Bounding Box

Fish bounding boxes were calculated for all 7, 237 images in which a fish was found. All but 263 of these were generated via pixel analysis, with the 263 falling back to the original `detectron` bounding box. 100 randomly-chosen images were reviewed manually to evaluate the calculation, a representative sample are presented in Figure 8. All 100 masks and bounding boxes were correctly placed on/around the location of the fish. However, a number of them lacked portions of lightly colored tails and/or fins. Specifically, 41 masks and bounding boxes covered the entire fish, 36 missed some of the tail, and 23 missed most or all of the tail, as seen in Figure 9. Masks and bounding boxes contain the head and trunk of the fish in nearly all cases, but further refinement of our algorithms will be needed to ensure that light fins and tails are masked consistently and accurately.

**Fig. 8.**
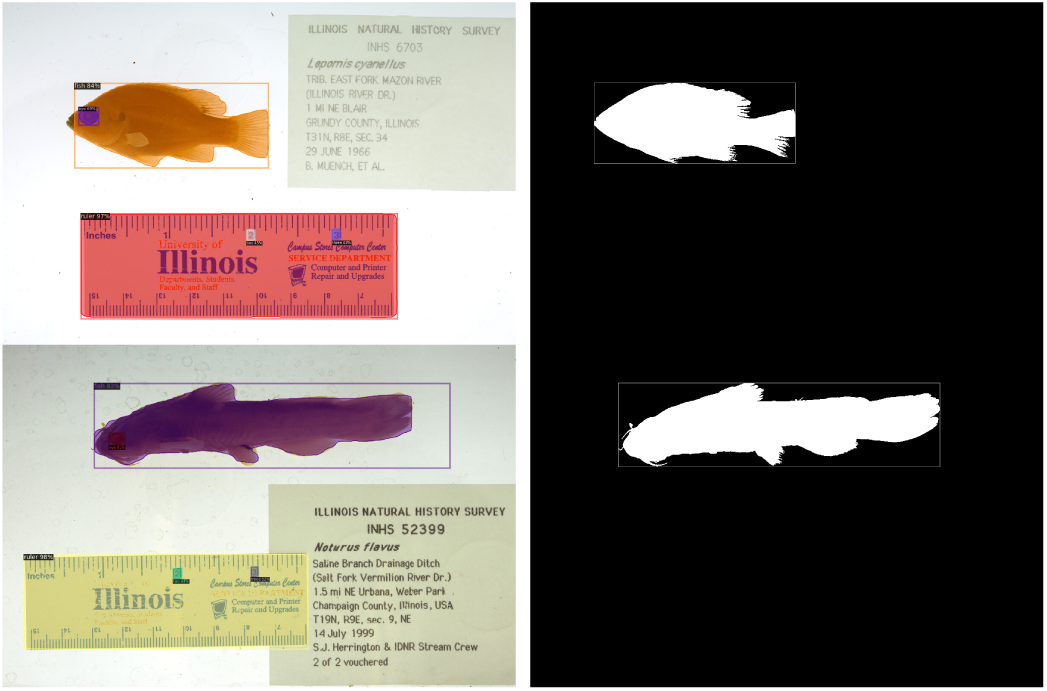
Examples of masks and bounding boxes from detectron (left) and pixel analysis (right).

**Fig. 9.**
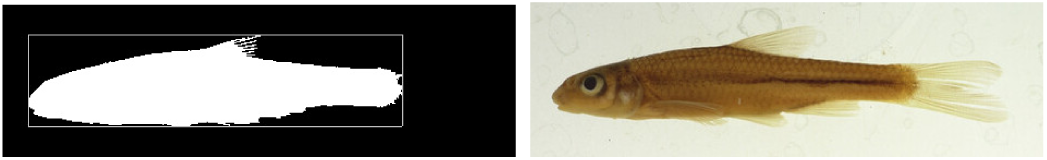
An example of a light colored tail being missed during the pixel analysis process.

### H. Scale and Length

Image scale and fish lengths were calculated for 7, 179 of the images. For the remaining 65 images, either the fish, the “2” and/or the “3” on the ruler were not detected. Image scale 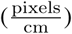 and fish length were measured, using ImageJ [41], in the same 100 test images. In this subset of images the average error for the scale calculation was 0.89%, and the average error for the fish length calculation was 5.55%. Scale calculations using the “2” and “3” method are nearly identical to those calculated by hand between the tick marks on the ruler. When the tail of the fish is accurately masked and the specimen is fairly straight, the length calculation is highly accurate as well. An example of such a result can be seen in Figure 10, for which the difference between the hand measured length and the automatically calculated length was only 0.6% (8.88 cm vs 8.82 cm). Thus, the primary means of lowering the error of the length calculation is to improve tail masking accuracy.

**Fig. 10.**
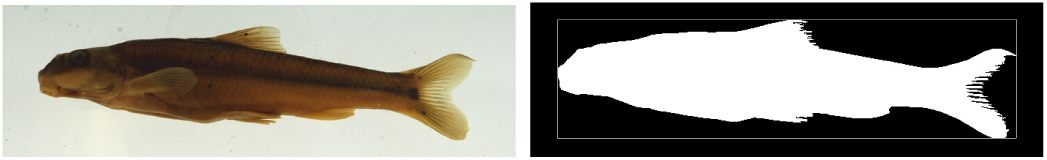
A fish for which the masking and measurement process was highly accurate.

## VI. Discussion

Overall our results show a proof of concept and offer a path forward for using object detection technology, enhanced by image informatics techniques, to improve and enrich metadata that enables advanced specimen image analysis and investigations of scientific research questions on an unprecedented scale. Our investigation has thus far focused on fish as the specimen of study. Fish are vertebrate animals (phylum Chordata), with over 34, 000 known unique species [42], with many more likely undiscovered. Species names are merely labels, and the discovery of species variation depends on both genotype and phenotype information. The ability to computationally analyze thousands of images of a single fish species, from different habitats and time periods, can lead to new discoveries that are impossible to pursue with manual methods. Digital library researchers have been concerned with computationally extracting image features, using content-based image retrieval methods. The work by Toress [43], while over 15 years old, demonstrates the challenges and opportunities to automatically generating useful metadata. Efforts to integrate such automatic metadata generation methods into digital library workflows and architectures still seem limited. This is likely due to the diversity of image shapes, sizes and the inconsistent configurations of specimens, labels, rulers, etc. within them. Object detection as explored in our research, working with an established architecture, is applicable to the larger world of biodiversity, well beyond fish, to include other fauna and flora, art and artefacts, and other digitized objects made accessible for scientific and scholarly research. Following object detection, one can apply pixel analysis and informatics methods to compute many more higher order properties from the initial segmentations.

Digital libraries serve to collect, provide access to, and archive rich collections of a wide array of materials. Many digital libraries interconnect with open repositories, supporting FAIR (Findable, Accessible, Interoperable, and Reuseable) [44]. In discussing the future of digital library services, Fox [45] underscores the need to prepare for and test ML applications. The work presented here, within the context of the BGNN project, demonstrates a clear need for improved metadata associated with specimen image collections. Much effort, time and money have been put into photographing and digitizing physical specimens, but without detailed and complete metadata properties the utility of these repositories for advanced computational analysis and ML is limited. Since it is very time intensive to generate all the pertinent properties by hand, automated techniques are essential to generating the missing metadata at scale.

## VII. Conclusion

In this paper we presented an automatic metadata generation approach. Using ML and image informatics algorithms, it is able to locate, mask and analyze specimens (currently limited to fish) in collection images with a high degree of accuracy. It produces 6 of the 22 core BGNN metadata properties [9], as well as image contrast, bounding boxes, scale and length information. Testing this approach on 7, 244 images from the INHS dataset [4], we see that the vast majority of the resulting metadata is correct within a tolerance of a few percentage points, and in some cases contains fewer mistakes than the manually generated validation data. Through further refinement and generalization beyond only INHS images, we aim to create a tool that can be distributed to specimen image collection curators to correct the metadata sparsity that precipitated this work.

### A. Future Work

The most pressing next step is to refine the pixel analysis thresholding process so that the entirety of even light colored fish are marked as foreground in the mask. A deficiency of the current process is that it only operates on single channel intensity. Some of the lightest tails appear yellow in hue to the human eye and easily distinguishable, but when compressed to a single intensity value they are almost identical in value to the white background. Considering when the RGB channels of a pixel are not equal in value may improve masking of such features. Another possible approach to solving this problem is to threshold and mask on subsets of the bounding box, as to ensure that very dark trunk pixels do not affect the thresholding of lighter regions.

Our longer-term goal is to create a generalized process that works on classes of specimen images. For the BGNN project we are beginning with fish images, but we are designing the metadata generation system so that it can eventually operate on other species if appropriately trained. To accomplish this, a much larger training dataset consisting of more diverse images will be required. The first step towards greater generality will be to operate on other fish collections besides INHS, which is something our program has already shown itself capable of doing during initial testing. Another requirement will be to generalize the ruler reading process beyond the INHS-specific reading of digits on the ruler, which will likely involve an automated method of reading ruler ticks instead of digits. Overall, the research reported in this paper will improve our BGNN workflow, and at the same time demonstrates an innovative approach that may greatly enhance digital library services for the tens of thousands of digitized specimens and images for other types of objects.

## VIII. Acknowledgment

We thank the full BGNN team for support, the data curation team at the Tulane University Biodiversity Research Institute, and Chris A. Taylor, Curator of Fishes and Crustaceans at the Illinois Natural History Survey (INHS). INHS is one of six fish collections participating in the Great Lakes Invasives Network (GLIN).

